# SAMBA: Structure-Learning of Aquaculture Microbiomes using a Bayesian Approach

**DOI:** 10.1101/2022.12.30.522281

**Authors:** Beatriz Soriano, Ahmed Ibrahem Hafez, Fernando Naya-Català, Federico Moroni, Roxana Andreea Moldovan, Socorro Toxqui-Rodríguez, M. Carla Piazzon, Vicente Arnau, Carlos Llorens, Jaume Pérez-Sánchez

**Author notes:** These authors contributed equally to this work (Beatriz Soriano, Ahmed Ibrahem Hafez).

## Abstract

In aquaculture systems, microbiomes of farmed fishes may contain thousands of bacterial taxa that establish complex networks of interactions among each other and among the host and the environment. Gut microbiomes in many fish species consist of thousands of bacterial taxa that interact among each other, their environment, and the host. These complex networks of interactions are regulated by a diverse range of factors, yet little is known about the hierarchy of these interactions. Here, we introduce SAMBA (Structure-Learning of Aquaculture Microbiomes using a Bayesian Approach), a computational tool that uses a unified Bayesian network approach to model the network structure of fish gut microbiomes and their interactions with biotic and abiotic variables associated with typical aquaculture systems. SAMBA accepts input data on microbial abundance from 16S rRNA amplicons as well as continuous and categorical information from distinct farming conditions. From this, SAMBA can create and train a network model scenario that can be used to: i) infer information how specific farming conditions influence the diversity of the gut microbiome or pan-microbiome, and ii) predict how the diversity and functional profile of that microbiome would change under other experimental variables. SAMBA also allows the user to visualize, manage, edit, and export the acyclic graph of the modelled network. Our study presents examples and test results of bayesian network scenarios created by SAMBA using data from: a) a microbial synthetic experiment; and b) the pan-microbiome of the gilthead sea bream (*Sparus aurata*) under different experimental feeding trials. It is worth noting that the usage of SAMBA is not limited to aquaculture systems and can be used for modelling microbiome-host network relationships in any vertebrate organism, including humans, in any system and/or ecosystem.

## 1. Introduction

Gut microbiomes in fishes and other vertebrates are subjected to complex and dynamic fluctuations that are driven by several factors associated with the host (e.g. genotype, physiological status, pathobiology) and its environment, lifestyle and diet [1]. In turn, each of these factors can contributing to improve the sustainability of industrial aquaculture [2]. Modelling complex relationships between the physical and biological components of aquacultural systems in the context of climate change and human population growth is one of the key future challenges in food production [3,4]. Current research on gilthead sea bream (*Sparus aurata*), a species frequently used in aquaculture in the Mediterranean, indicated that the gut microbiota is a reliable criterion to evaluate the success of selective breeding with changes in diet composition [5-7]. However, our understanding about these kinds of dynamics is at an infancy state, due to their inherent complexity, the multiple biotic and abiotic factors involved, and the enormous variability of mucosal microbial populations among distinct individuals of the same population [8,9]. For this reason, there is a growing need to develop tools that can model how fish microbiomes and their hosts interact under variable farming conditions. Aling these lines, bayesian networks (BN) and structure learning [10-12] may be especially useful due to their capacity to infer directional relationships in microbial communities [13,14].

BNs are probabilistic graphical models based on Bayes Theorem that represent and evaluate the conditional dependencies among a set of variables via directed acyclic graphs (DAG): variables and their interrelations of dependency are represented as nodes and edges respectively [15]. On the other hand, structure learning refers to the process to learn the structure of the DAG from the available data, creating a model where an edge between two nodes indicates direct stochastic dependency, while no connection (edge) between two nodes identifies that the corresponding variables are independent or conditionally independent [13]. BNs have been used to improve our knowledge of sustainable aquacultural systems [16]; however, they are yet to be applied to reveal dynamic interactions between different biotic and abiotic factors in aquacultural systems. Moreover, BN tools are typically tailor-made solutions created using command line interface (CLI) software packages (e.g. bnlearn). As such, they are complex to manage and often only practical tools for expert bioinformaticians and computational biologists [17]. Indeed, most user-friendly BN tools with Graphical User Interfaces (GUI), such as ShinyBN [18] or BayesiaLab [19], only work with discrete variables and small datasets. In addition, while the recently released BayeSuites tool [11] manages continuous variables and large datasets, it currently cannot make inferences and establish conditional probability distributions based on discrete variables. To address these issued, we created SAMBA (Structure-Learning of Aquaculture Microbiomes using a Bayesian Approach).

SAMBA is a new BN tool to investigate microbiome-systems or microbiome-host network dynamics in aquaculture systems by modelling how fish gut microbiomes and/or pan-microbiomes interact with the various biotic and abiotic factors. Here, we provide examples of the functionality of the tool’s web-interface and evaluate how SAMBA performance when building and estimating the conditional dependencies among the variables in the DAG model. To this end, we will use two training datasets of different nature and complexity: i) artificial microbial community with few taxa and defined composition; and ii) real fish microbial communities of S. aurata resulting from a given time and aquaculture infrastructure with diet as the main changing experimental variable.

## 2. Material and Methods

### 2.1. SAMBA Modules

SAMBA is currently available in a GitHub public repository. The URL for downloading the input data is reported in the Data Availability Statement. The tool can be installed on personal computers, and it is based on a backend engine core that consists of a set of workflows and pipelines implemented in R and Python using third-party software dependencies. The frontend component of SAMBA consists of a web-based Graphical User Interface (GUI) implemented using shiny [20] to provide a friendly and intuitive interface to manage the engine core. Functions and tasks of SAMBA are structured in five modules: “Build”, “Inference”, “Prediction”, “Viewer”, and “Downloads”. A User Guide is provided in Supplementary file 1.

#### “Build”

This module creates and trains BN models from the provided input data using a pipeline based on the bnlearn R package [17]. This module works with continuous and discrete variables. However, a discretization optional step is implemented using one of the following methods: interval, quantile or Hartemink [21]. For those continuous variables that are not discretized, a Shapiro [22] test is performed to know if they follow a normal distribution. For further information, please refer to Supplementary file 1. To create the BN models, SAMBA allows the user to fit distribution parameters. The current implementation includes a normal distribution in a logarithmic scale (Log-Normal) and a generalized linear model, the Zero-inflated Negative Binomial (ZINB) distribution, that better fits highly dispersed data with an excess of zeros in the taxa abundance counts [23]. The hc() function of bnlearn learns the structure of a BN using a hill-climbing greedy search (score-based algorithms). According to Scutari et al. 2019 [24], these algorithms are usually faster and more accurate for both small and large sample sizes. The hill-climbing search [25] explores the space of the DAG by single-arc addition, removal, or reversals. It also assigns a rate to the BN model using catching, decomposability, and three equivalence score functions to reduce the number of duplicated tests [17]. The three score functions are: Akaike Information Criterion (AIC), Bayesian Information Criterion (BIC), and Multinomial log-likelihood (loglik). The score method is adjusted for the hc() method so that a higher score is generally preferred.

The training of the model constructed by the “Build” module is performed by using the bn.fit() function of bnlearn or the aforesaid bn.fit() combined with the zeroinfl() function from the pscl package [26]. The bn.fit() function fits, assigns, or replaces the parameters of a BN conditional on its structure, while zeroinfl() fits zero-inflated regression models for count data via maximum likelihood. Once the parameters have been fitted, the strength of each connection is calculated using BIC and Mutual Information (MI) criterion [27] and the arc.strength() function of bnlearn to remove all links with strength values greater than the user-defined threshold. The future() function of the future package [28] allows the user to continue using other functions in the app while a model is being computed.

#### “Inference”

This interface uses functions of bnlearn and dagitty [29] to infer how the diversity of the pan-microbiome indexed in the BN is influenced by the experimental variables (season, diet composition, temperature, genetics, etc.). The “Inference” module provides two different report options: Conditional probability tables (CPTs) and DAG. The CPT option uses a cosponsoring quantile-quantile plot of the fitted node to show the type of relationships among different taxa under different experimental variables. The DAG option creates a DAG from the bnlearn output using the dagitty() method of dagitty and allows the markov blanket of a given node to be extracted from the DAG using a markovBlanket() method.

#### “Prediction”

This interface allows the user to manage two predictive pipelines. The first, “Predict abundances” is a workflow based on bnlearn and pscl that predicts how taxa abundance counts will likely change based on changes in one or more farming variables selected as conditional evidence. In the Log-Normal Distribution mode, normalized frequencies in log scale are obtained via the cpdist() function of bnlearn. In the ZINB distribution mode, a custom sampling method from the fitted ZINB models of each taxon in the BN. The second pipeline of this module, “Predict Metagenomes”, infers the metagenome of a given pan-microbiome under specific experimental variables. It uses PICRUSt2 [30], which includes two different database annotation protocols: MetaCyc [31] and KEGG [32]. For more details about the PICRUSt2 workflow and its dependencies [33-37] please refer Supplementary file 1.

#### “Viewer”

This provides the user with tools to visualize, edit, customize, navigate, and export the DAG in various graphical formats. The “Viewer” is implemented using commands from several different packages. The subgraph() function of bnlearn plots the graph. VisIgraph() and the renderVisNetwork() functions of visNetwork [38] both provide an interactive display of the DAG. The strength.viewer() function of bnviewer [39] shows the strength of the probabilistic relationships of the BN nodes. It also uses model averaging to build a graph containing only significant links. The decompose() function of the igraph package [40] visualized specific node groups and thus allows users to work with a specific subgraph. CPTs with conditional probability information about the inter-relations of each node can be displayed in the viewer using the datatable() and dataTableProxy() functions of the DT package [41]. These functions allow the user to browse and filter information from the DAG. The “Viewer” also integrates a sidebar with tools for highlighting, selecting, and/or editing specific nodes and features using functions from the VisNetwork package, such as visOptions(), visRedraw(), visSetSelection(), visUpdate-Nodes(), and visUpdateEdges(). It also contains JavaScript code introduced through the runjs() function of the shiny package [42] and the JS() function of the htmlwidgets package [43]. Graphs can be downloaded as HTML, PNG, JPEG, or PDF files. A screenshot can be taken of the current network and exported using the shinyscreenshot package [44].

#### “Downloads”

This is a repository for the user to download .zip files containing results and output files from the “Build” module (the output includes normalized counts, link strengths, and a RData file containing the BN model) or the “Prediction” module (provided by metagenomic prediction).

### 2.2. Industrial testing dataset (sequencing and dataset definition)

Semi-synthetic bacterial community ZymoBIOMICS™ Microbial Community Standard II (Zymo Research Corp., CA, United States). This “Mock community” is composed by seven bacteria (*Listeria monocytogenes, Pseudomonas aeruginosa, Bacillus subtilis, Salmonella enterica, Escherichia coli, Lactobacillus fermentum and Enterococcus faecalis*) with known differential abundances, distributed on a log scale, ranging from 0.0001% (*E. faecalis*) to 95.9% (*L. monocytogenes*). Eight replicates of the Mock community were sequenced for 16S rRNA genes using the Oxford Nanopore MinION platform under two PCR conditions: PCR1 (Temperature of annealing 55_O_C and 25 PCR cycles) and PCR2 (Temperature of annealing 52_O_C and 30 PCR cycles) [45]. The raw abundance counts and the PCR conditions to sequence the mock community were used as input to SAMBA. Full details about this dataset are provided in Supplementary file 2 and the URL for downloading the input data is reported in the Data Availability Statement.

### 2.3. Empirical testing *S. aurata* dataset (sequencing, experimental design, rearing conditions, and dataset definition)

Intestinal pan-microbiome data were taken from the results of three published experimental trials that used S. aurata as case study host model [46-48]. The *S. aurata* dataset includes 844 taxa classified at the genus level and obtained by sequencing the V3-V4 hypervariable regions of 16S rRNA. The autochthonous microbiota populations were sequenced from both anterior and/or posterior intestine sections of 72 randomly selected specimens of *S. aurata*. Briefly, the trials were conducted in parallel (Spring-Summer 2020) at the Institute of Aquaculture Torre de la Sal under natural light and temperature conditions (40°5′N; 0°10′E) using fish with the same genetic background (sibling animals from the same hatchery batch; Avramar, Burriana, Spain). The resulting microbial populations were investigated in relation to different feeding scenarios (LSAQUA, EGGHYDRO, and GAIN_PRE_ that are summarized in Table 1 In the LSAQUA trial, fish meal (FM) was either partially (50%) or completely (100%) substituted with a protein replacer of processed animal proteins (PAPs) and bacterial single-cell proteins (SCPs). In the EGGHYDRO trail, combinations of FM and fish oil (FO) were used with or without a bioactive egg white hydrolysate. In the GAIN_PRE trail, FM was completely replaced with alternative protein sources (aquaculture by-product meal, insect meal, microbial biomass, and plant proteins) supplemented with a commercially available health-promoting feed additive. The performance of SAMBA was tested using the S. aurata dataset to build a BN model under the following parameters: BN score (BIC), taxa distribution (ZINB), link strength thresholds (MI<0.05; BIC<0), and a prevalence filter of 50 (which dismiss those taxa that are present in less than 50% of the samples with at least a minimum abundance of 1 read count). The experimental variables were considered independent so that predictions could be made for scenario that were not directly tested in the methodology. The predictive performance of SAMBA was evaluated in the EGGHYDRO trial. In addition, we evaluated how the predictive function of SAMBA to forecast how the most likely distribution of pan-microbiome abundances can change in response to modification in one of the variables. Specifically, it was tested how modifying the modification of FM variable would affect the output of EGGHYDRO trial scenario 3. This new Scenario, designated as Scenario 4, was designed with these experimental variables: FM >20; FO ≤ 4; “AI” Tissue; “EWH” additive.

**Table 1.**
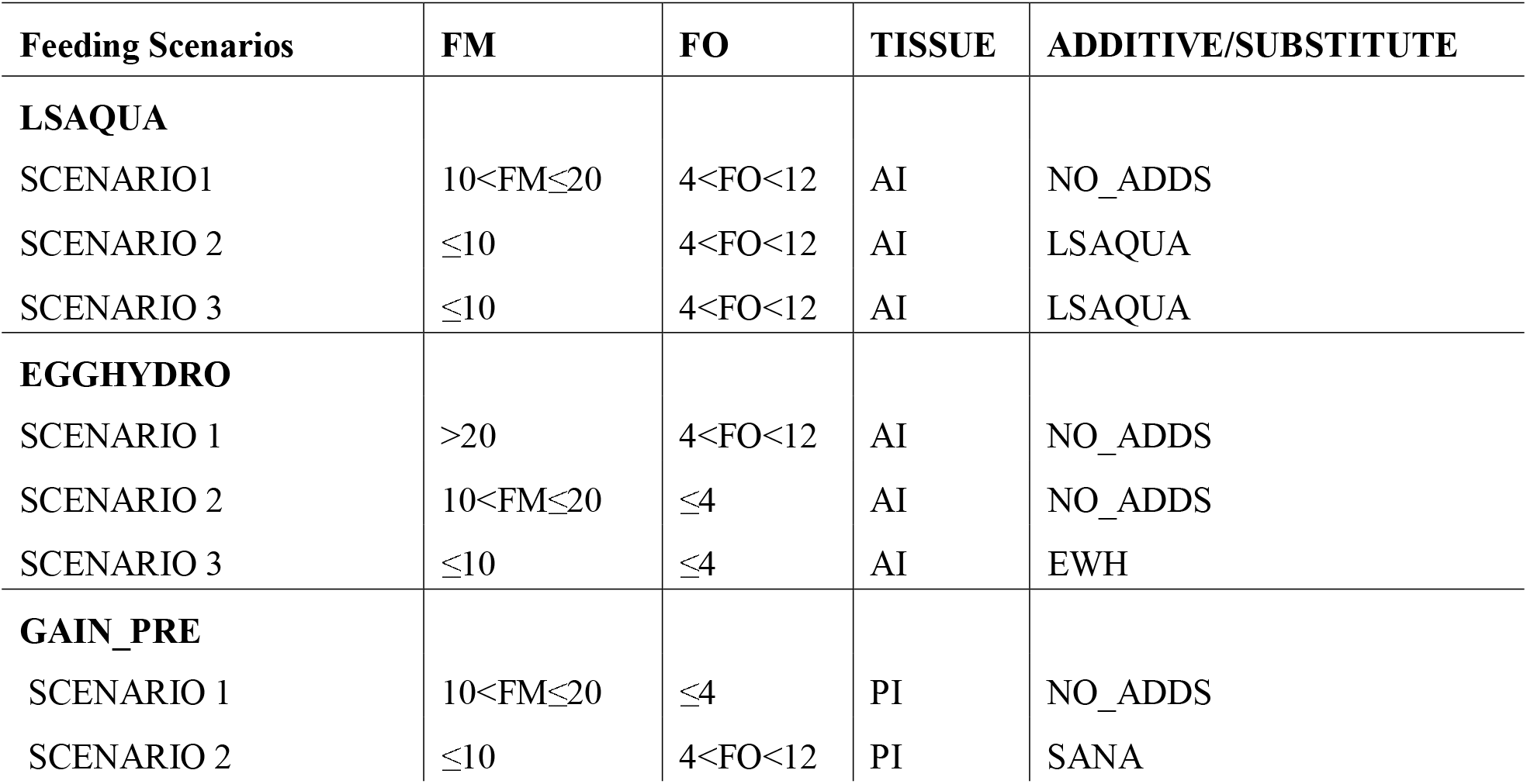
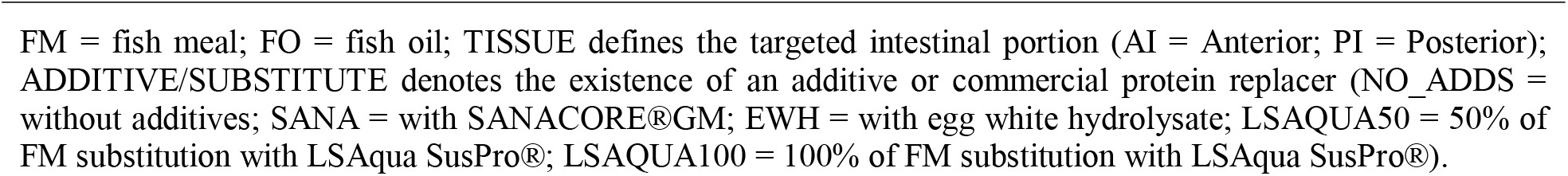
Experimental variables for the three Gilthead Sea Bream farming trials

| Feeding Scenarios | FM | FO | TISSUE | ADDITIVE/SUBSTITUTE |
| --- | --- | --- | --- | --- |
| **LSAQUA** |  |  |  |  |
| SCENARIO1 | $10 < FM \leq 20$ | $4 < FO < 12$ | AI | NO_ADDS |
| SCENARIO 2 | $\leq 10$ | $4 < FO < 12$ | AI | LSAQUA |
| SCENARIO 3 | $\leq 10$ | $4 < FO < 12$ | AI | LSAQUA |
| **EGGHYDRO** |  |  |  |  |
| SCENARIO 1 | $> 20$ | $4 < FO < 12$ | AI | NO_ADDS |
| SCENARIO 2 | $10 < FM \leq 20$ | $\leq 4$ | AI | NO_ADDS |
| SCENARIO 3 | $\leq 10$ | $\leq 4$ | AI | EWH |
| **GAIN_PRE** |  |  |  |  |
| SCENARIO 1 | $10 < FM \leq 20$ | $\leq 4$ | PI | NO_ADDS |
| SCENARIO 2 | $\leq 10$ | $4 < FO < 12$ | PI | SANA |

FM = fish meal; FO = fish oil; TISSUE defines the targeted intestinal portion (AI = Anterior; PI = Posterior); ADDITIVE/SUBSTITUTE denotes the existence of an additive or commercial protein replacer (NO\_ADDS = without additives; SANA = with SANACORE®GM; EWH = with egg white hydrolysate; LSAQUA50 = 50% of FM substitution with LSAqua SusPro®; LSAQUA100 = 100% of FM substitution with LSAqua SusPro®).

## 3. Results

In this article, we introduce SAMBA, the software implementation of a BN approach to learn, build, and train BN models from input datasets with quantitative and qualitative variables (including taxa abundance raw counts). As shown in Figure 1, SAMBA has a user-friendly GUI interface that provides access to five modules (“Build”, “Inference”, “Prediction”, “Viewer” and “Downloads”), which overcome the usual computational complexity that exists in the modelling of BNs (see Supplementary file 1 for technical details). The application can be used to investigate the causal relationships between microbiomes and their hosts by deciphering how the taxa population are related each to other and influenced by the experimental variables. SAMBA can also be used to navigate the built BN model and to inspect the distribution of conditional dependences among the distinct variables, identifying those that provide statistically significant information about how a change in feed formulation, or any other environmental condition, may derive in a modulatory effect in the microbial profile. Additionally, SAMBA conditional BN dependencies provide a system biology perspective that, in comparison to conventional analyses based on the relative abundance of the taxonomic groups, is very informative because the user may find combined actions between taxa without making independence assumptions, as normally done in a usual 16S count analysis.

**Figure 1.**
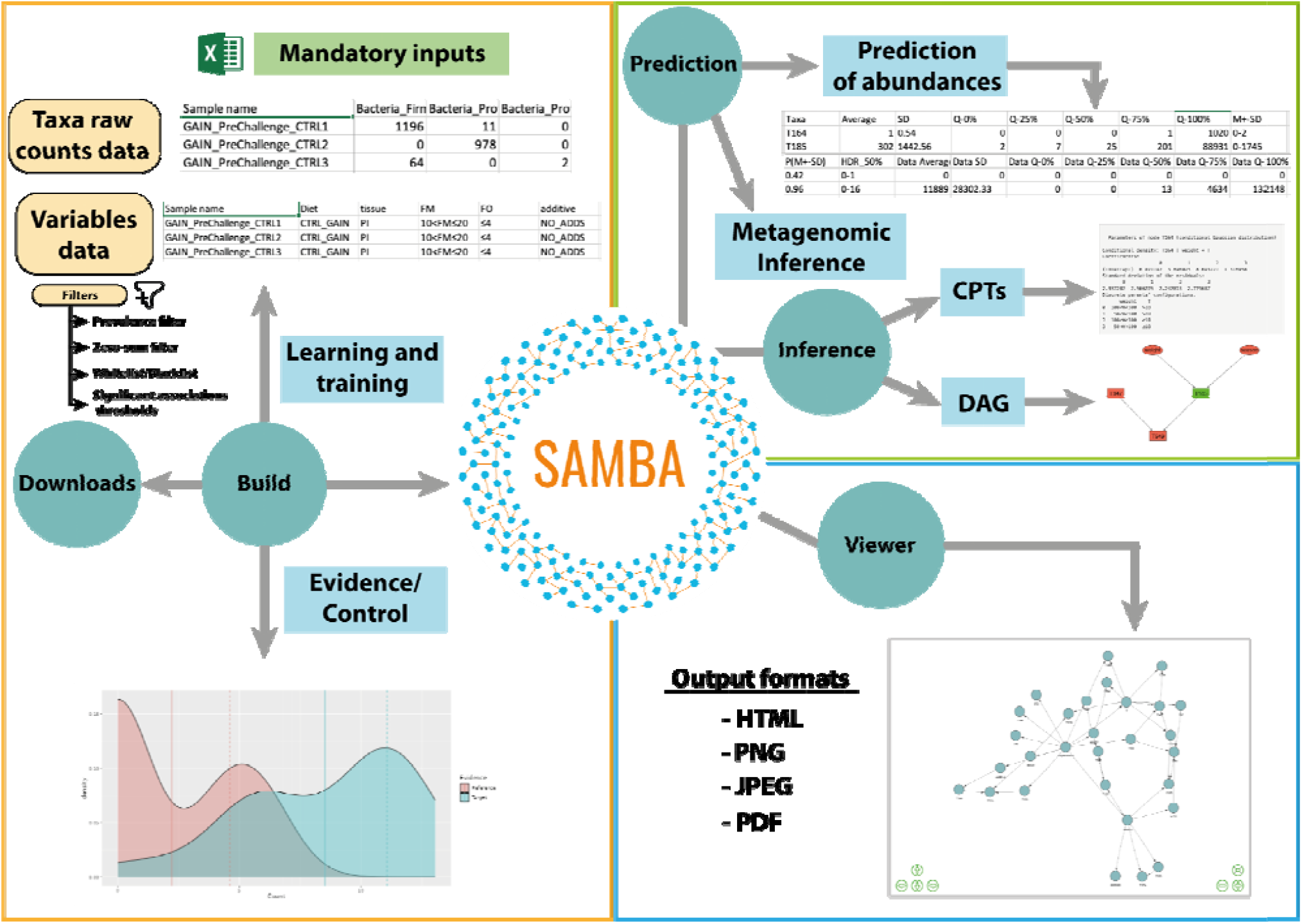
Graphical description of the functions and interface panels in each module of SAMBA. Green circles represent modules and blue squares represent interfaces.

To test the potential and predictive power of SAMBA, we fed the tool with two training datasets (the Mock community and the *S. aurata* dataset) to create two BN models. First, we addressed the mock community because the simplicity nature of this dataset. The mock community model offers a controlled semi-synthetic scenario based on seven taxa with known abundances counts (i.e. with no natural inter-sample dispersion among the abundance counts of the modelled taxa) and one experimental variable (the PCR). In Figure 2A we show the DAG resulting from the mock community BN model, which depicts how the seven taxa of this community are connected each to other in the network as a consequence of their abundances and the experimental PCR condition (with the exception is *L. fermentum*, which is not affected by the PCR condition because is the least abundant taxon. Predictions based on the mock community BN model were also performed and provided in Supplementary file 2. In addition, Figures 2B and 2C are two correlation plots which show a remarkable linearity between the observed and the predicted abundances (under the two PCR conditions) of the seven taxa constituting mock community. These two analyses are both supported by correlation coefficients over 0,99 for both PCR1 and PCR2 conditions, and with p-values for a F-test of 6,12E-10 and 1.48E-22 in an analysis of variance (piece of information also available in Supplementary file 2). We can thus conclude that SAMBA predictions accurately approximate the real-world observation from labmade microbial simplified scenarios that could be particularly helpful in designing and exploring synthetic biology experiments.

**Figure 2.**
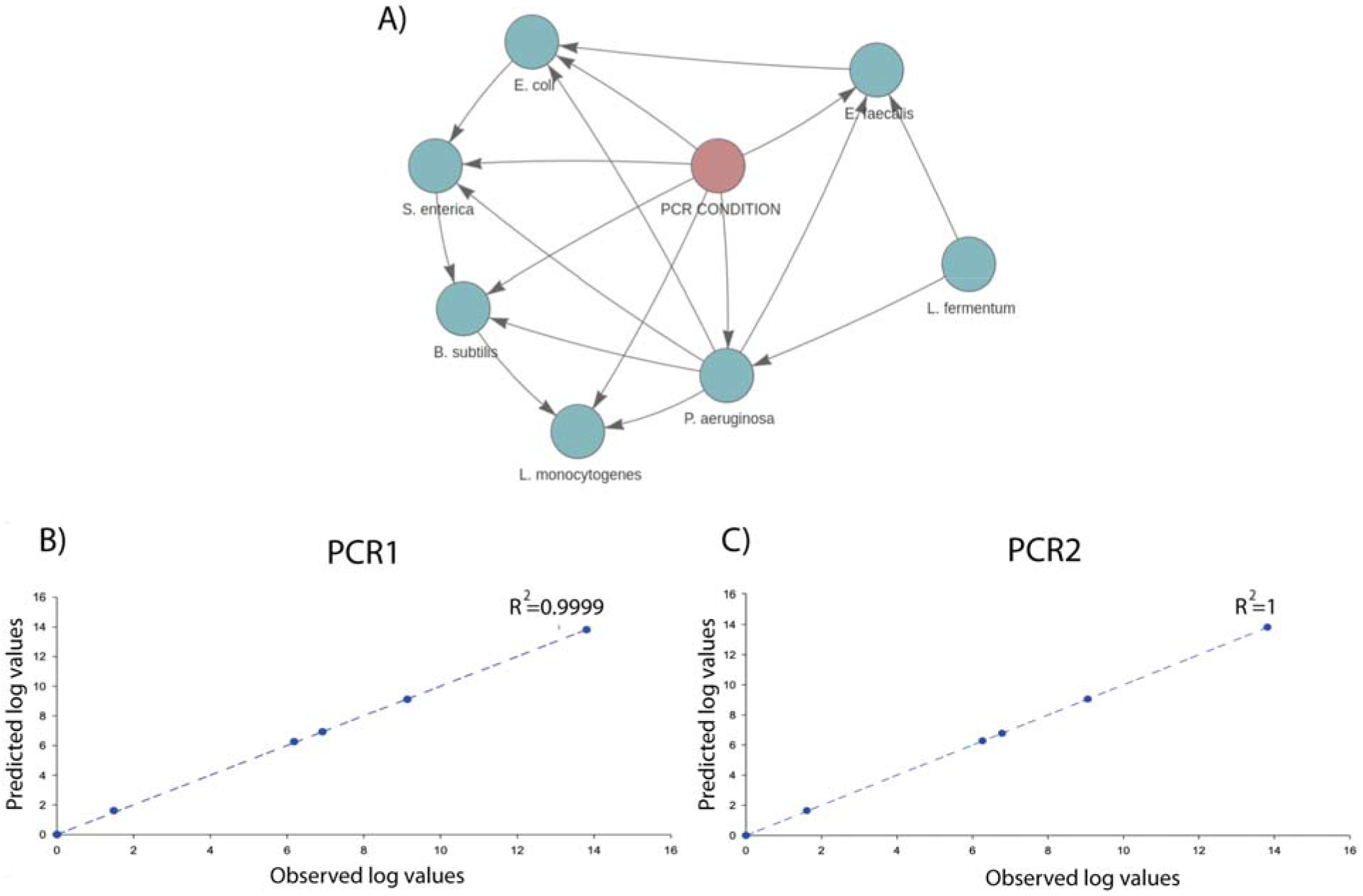
A) Screenshot of the BN model created by SAMBA for the mock community showing how the distinct taxa are related to each other; B) Correlation analysis between the average predictions for the seven taxa relative to the normalized average values of abundances, in log scale, under PCR1 condition; C) Same correlation analysis but under PCR2 condition

The second BN approach based on the *S. aurata* dataset was performed to assess how SAMBA builds and manages BN models using microbiome data from real-world experimentation (i.e. with high levels dispersion in the taxa’ abundance counts). In Figure 3A, we show the DAG of the BN built using the *S. aurata* dataset. The prevalence filter of 50% reduces the number of taxa to 45. This number represents the core microbiome present in at least 36 of the 72 samples. This is a useful feature of SAMBA as it allows the user to focus not only on the whole dataset or the most abundant OTUs in the bacterial populations (the usual approach of 16S metagenomic analyses), but also on different subsets by managing the filtration parameters of the interface. Extracting functional and quantitative information from the DAG with SAMBA is easy and intuitive with the inference module. In particular, an example of how these results can be obtained is reported in Figure 3B, with the Markov blanket graph extracted from the total DAG. The image shows that the node representing Pseudomonas is the child of the nodes representing the experimental variables FO, FM and *Phyllobacterium* that indirectly connects *Pseudomonas* with *Clostridium sensu stricto*. The coefficient for conditional probability distribution that is significant for the node *Pseudomonas*, and the results of a Z test for each coefficient are shown in Supplementary file 3. According to this, it is possible to observe in Figure 3B that the FM variable (in its three states) has a significant impact on the abundance of Pseudomonas. In contrast, the variable FO only significantly impacts on *Pseudomonas* when FO≤4, but not when 4<FO<12. Additional to the dependences that occur between experimental variables and taxa, the Markov blanket also offers information regarding the taxa-taxa interaction. The presence of *Phyllobacterium* in the pan-microbiome is significant in respect to *Pseudomonas*. This relationship means that the abundance of *Phyllobacterium* influences abundance of *Pseudomonas*. Nevertheless, the impact of *Phyllobacterium* is less than that of FM and FO, which are the main causes for the variability in abundance of *Pseudomonas* in the *S. aurata* BN model.

**Figure 3.**
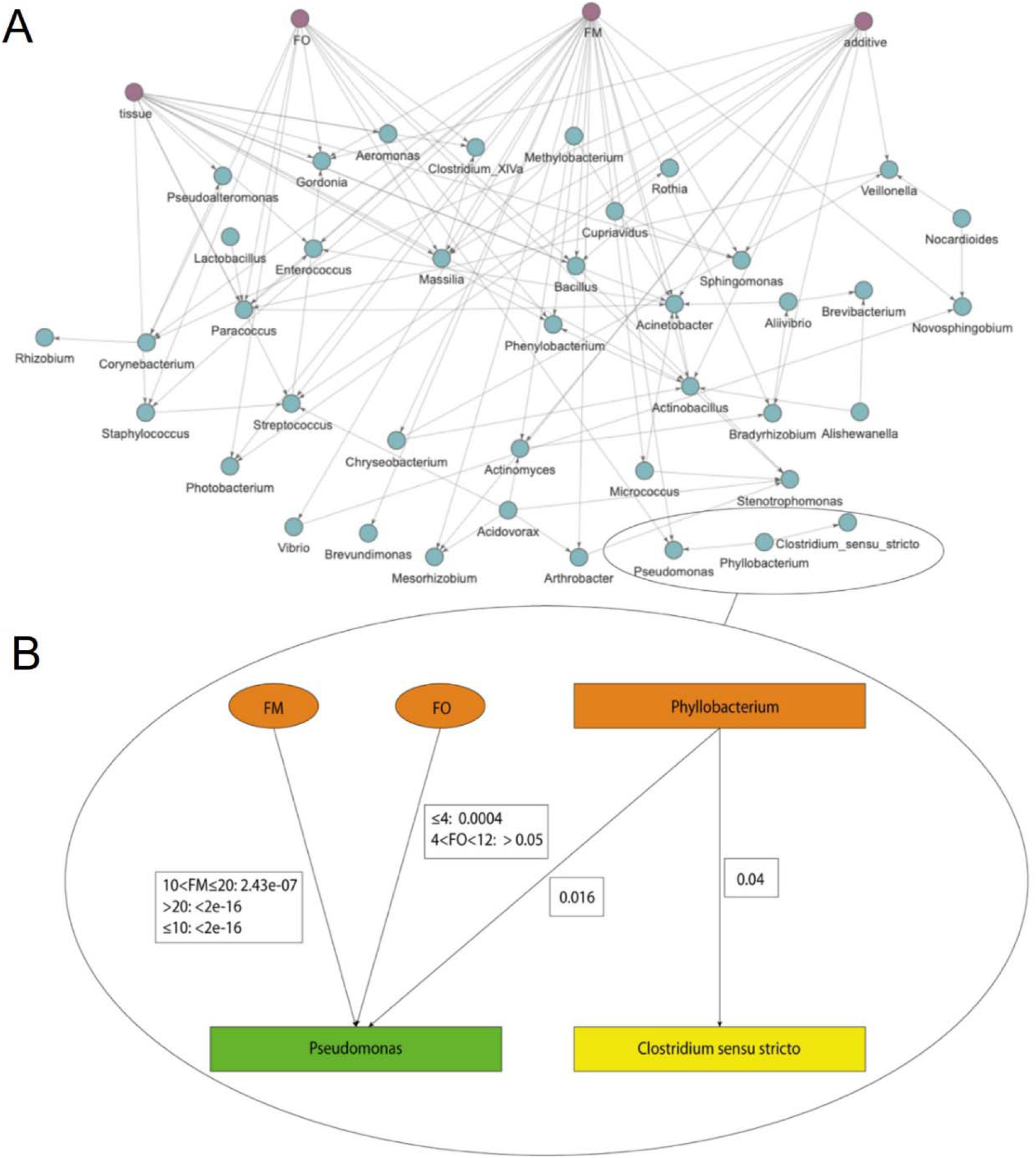
A) *S. aurata* model DAG built with SAMBA using the ZINB distribution showing the significant edges calculated between the experimental variables. Experimental variables are in pink while taxa are blue. B) Markov blanket of *Pseudomonas* node extracted from the total DAG. Taxa are represented with rectangles while experimental variables are represented with ovals. The Pseudomonas node is green, nodes with direct relationships with *Pseudomonas* (FM, FO and P*hyllobacterium*) are orange, and nodes with indirect relationships are yellow. The p values refer to whether a taxon or a variable has a significant influence (p, 0.05) on the *Pseudomonas* node.

The *S. aurata* dataset was also used to test the predictive capability of SAMBA. The profile of taxa abundances was predicted for the three feeding scenarios of the EGGHYDRO trial (Scenarios 1, 2 and 3 in table 1) using the distribution of probabilities provided by the *S. aurata* BN model. Full predictive reports for Scenarios 1,2 and 3 are available in Supplementary file 4, which shows that the probability density value of the range of the generated samples (the (P(M+SD)) was above 0.60 for 44 of the 45 taxa in Scenario 1; for 39 of the 45 taxa in Scenario 2; and for 42 of the 45 taxa in Scenario 3. This means that under Scenarios 1, 2 and 3 the ranges of predicted values made by SAMBA is significant for 98%, 87%, and 93% of the taxa, respectively. Moreover, with some reasonable exceptions, the mean, median, standard deviation, and quantiles overlaps with the SAMBA predictions (Figures 4, and 5). Thus, we conclude that SAMBA accurately predicts taxa abundances in a large matrix of data, even if their abundance distribution show significant dispersion due to the inter-sample biological variability, like real microbiota data [8,9].

**Figure 4.**
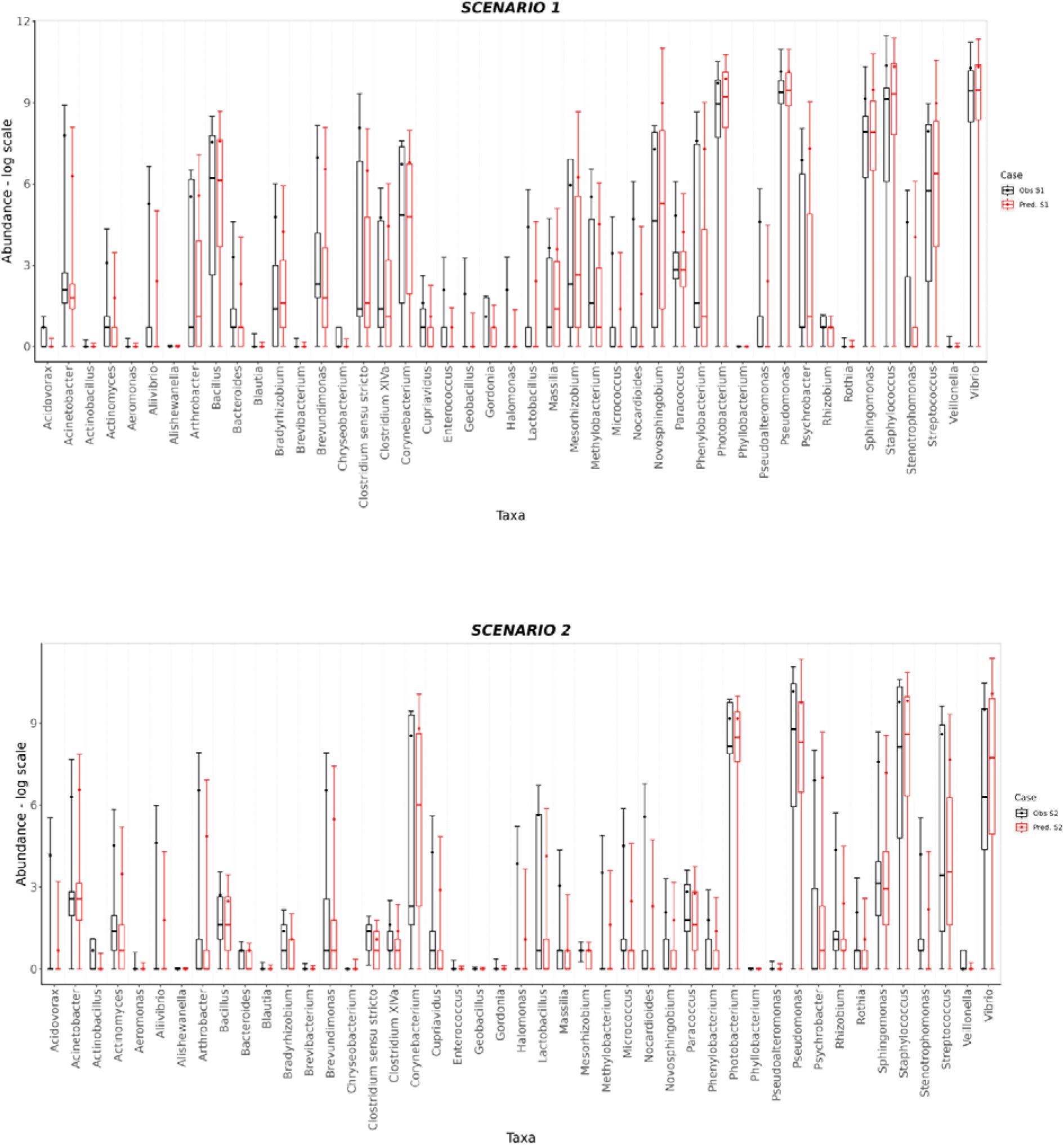
BoxPlot comparisons between predictions (red) and experimental observations (black) in log scale for Scenario 1 and Scenario 2. In both cases, the box plot for each taxon covers a range of values defined by the average abundance for that taxon and its standard deviations. 25 and 75 % quantiles as well as the median abundance values are represented as boxes.

**Figure 5.**
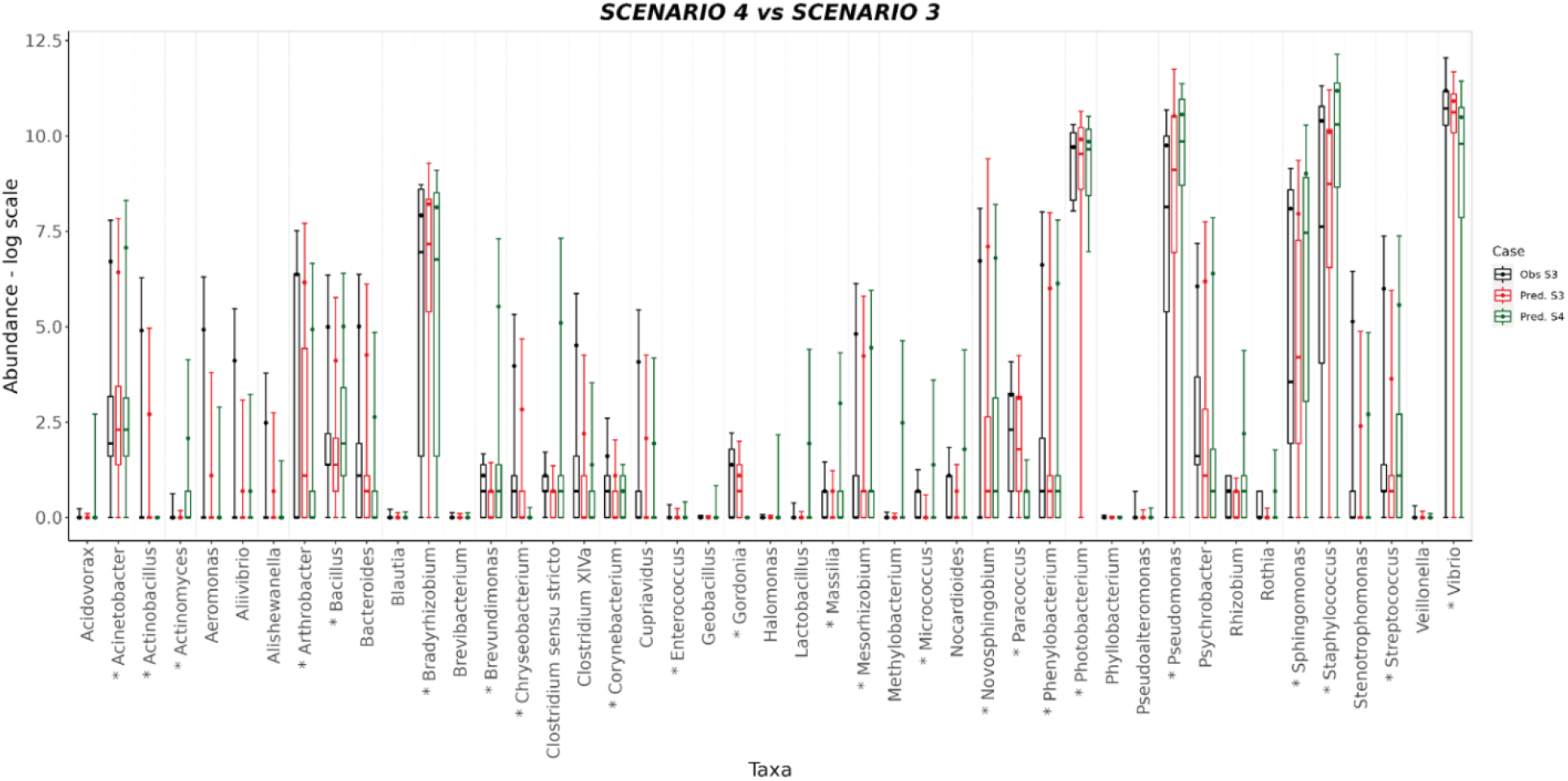
BoxPlot comparisons between predictions and experimental observations in log scale for Scenario 3 and Scenario 4 which is a prediction about how Scenario 3 would likely change when changing the FM conditions. Predictions for Scenario 3 are represented in red and observations black. Predictions for Scenario 4 are represented green. No observed data is provided for Scenario 4 because it is a virtual scenario that derives from combining experimental condition FM of Scenario 1 with the experimental conditions for FO, TISSUE, and ADDITIVE of Scenario 3. As in Figure 4, each box plot includes information from average abundance and standard deviations for each taxon plus the quantiles 25 and 75 and median for the distribution of abundance counts. Taxa that are significantly influenced by variable FM are highlighted with an asterisk (see also Figure 3).

Scenario 4 also tested the predictive capability of SAMBA by combining information from two different trials. More specifically, Scenario 4 predicted how the profile of taxa abundances of Scenario 3 would likely change under the experimental condition of Scenario 1 when FM>20 (Supplementary file 4). In Figure 5, we also plotted mean, median, standard deviation, and quantiles obtained from Scenario 4 to compare with those from Scenario 3. The figure shows how the predicted abundance of some taxa significantly changed is response to FM. Thus, this suggests that SAMBA allowed virtual scenarios virtual scenarios to be explored by the BN model. This feature of SAMBA is useful for improving both productivity and sustainability of fish farms. This is because it can simulate, a priori, the effects in the microbiome-host interrelations under different conditions (e.g., new diet, additives, or a modification of the environmental conditions).

When considered together, our results highlight the capacity of SAMBA to identify which FM or FO dietary levels, have a significant influence on the microbiota profile of farmed fishes. Understanding these associations emphasizes the ability of SAMBA to predict changes in the microbial communities of *S. aurata* as a case study of aquatic farmed animals. In addition to this, the information that comes from taxa interactions can give interesting information on the dynamics of a collaborative and/or competitive nature that rule a complex environment such as the intestinal biome. Future use of SAMBA will expand the available testable experimental conditions, allow users to customize their analysis, and introduce their personal expertise and knowledge when modelling a microbe population. SAMBA can therefore be a valuable tool for making the research in the aquaculture field more dynamic.

The use of SAMBA is not restricted to fish farming and aquaculture and it can be adapted to build microbial host BN models from other systems and/or vertebrate organisms, including humans. In fact, the input dataset accepted by SAMBA only consists of two files: one with the abundance counts per bacterial taxa and another with the experimental/environmental variables. In the present case study, the experimental variables were discrete (ergo categorical). However, the tool is able to manage both continuous and categorical experimental variables, such as sex, age, specimen size, genetic background, tissue, season, diet composition, pH, temperature, phenotypes, and more. Furthermore, in this version of SAMBA, we used a single omics (16S amplicons) model to assess microbiome-host network interrelations. However, future updates will integrate multi-omics variables from RNAseq and Methylseq, as well as other layers of information. This will extend the applicability of SAMBA to other topics where BNs have been proven to be effective, such as behavioral and welfare assessments, epigenomics, and genomics and transcriptomics, [49-51].

Regarding data distribution, the tool is currently limited to two models (Log-normal and ZINB). Future versions will include other Generalized linear models such as Negative Binomial, Poisson, and Gamma [23,52]. Additionally, different normalization methods, including for instance scaling factors for data correction per sample depth and gene size will be considered (RPKM, FPKM, TPM, RMS, etc.) [53]. We also aim to implement tools for determining the minimum training sample size needed to detect a significant effect for a given experiment [54].

In summary, the first SAMBA release highlights how BN models are valid computational approaches to investigate host-microbiome aquacultures, and other systems. The tool deconstructs and quantifies the structure of network relationships influencing the microbial dynamics of a given aquaculture and from that point on permits the user to obtain realistic predictions not only from tested and for inferred experimental conditions. We are committed to escalate SAMBA with new functions, databases, and tools including the implementation of new layers of information to also address multi-omic approaches.

## Supplementary Materials

The following supporting information can be downloaded at: www.mdpi.com/xxx/s1, Supplementary file S1. SAMBA’s User Guide. The file contains a user guide indicating how to use the different modules that compose SAMBA. Supplementary file S2. Coefficient for conditional probability distribution for the node Pseudomonas extracted from the DAG represented in Figure 5 and Z test for significance. Supplementary file S3. Mock community BN model. Excel file with three tabs: Tab1) Summary of results and input data; Tab2) Regression analysis and F-test for ANOVA under PCR1 condition; Tab3) Regression analysis and F-test for ANOVA under PCR2 condition. Supplementary file S4. S. aurata BN model. Excel file with four tabs for the EGGHYDRO trial predictions; Tab1) Scenario 1 (under experimental variables FM >20; 4<FO<12; “AI” Tissue; “NO_Adds” additive); Tab2) Scenario 2 (10<FM≤20; FO ≤4; “AI” Tissue; “NO_ADDS” additive); Tab3) Scenario 3 (FM ≤10; FO ≤4; “AI” Tissue; “EWH” additive); Scenario 4 (FM >20; FO ≤4; “AI” Tissue; “EWH” additive).

## Author Contributions

Conceptualization, B.S., A.I.H, F.N-C., M.C.P, V.A., C.L. and J.P-S.; Methodology, B.S., A.I.H., F.N-C., C.L. and J.P-S.; Software Programming, B.S., A.I.H. and R.A.M.; Application testing, F.N-C., F.M. and S.T-R.; Manuals and tutorial resources, B.S. C.L. and R.A.M.; Writing and manuscript preparation, B.S., F.N-C., A.I.H., F.M., M.C.P., C.L. and J.P-S. All authors read and approved this work.

## Informed Consent Statement

Not applicable.

## Data Availability Statement

The source code of SAMBA as well as a dataset for testing the application are available at https://github.com/biotechvana/SAMBA. The two testing datasets (“Mock_community” and “S. aurata dataset”) used to evaluate the performance and accuracy of SAMBA as well as the R.Data file with their respective BN models are available at the Web site of SAMBA with the following URL address https://github.com/biotechvana/SAMBA/tree/main/Testings_datasets. Each dataset includes two files containing the experimental variables (i.e. diet, tissue, additive, etc.), and the raw counts of all taxa per amplicon sample. This URL also includes another dataset named “metagenome_testing_dataset” which is provided for users interested in training examples for metagenome predictions using the “Prediction” module of SAMBA. Original Fastq files from Mock community, the LSAQUA, EGGHYDRO, and GAIN_PRE trials are available at the SRA archive with the following bioproject accessions PRJNA891255; PRJNA713764; PRJNA705868; PRJNA750446.

## Acknowledgments

We thank Nathan J Robinson for critical reading and corrections. We also thank the reviewers of the Genes Journal for their criticisms and suggestions that helped us to improve our article.

