## Supplementary file 1 for "SAMBA: Structure-Learning of Aquaculture Microbiomes using a Bayesian Approach"

**Supplementary File 1: User Guide of SAMBA.**

In this section we provide a user guide tutorial explaining how to manage the distinct sections of SAMBA. Most of the functions described in this section are provided by one of the following R packages: *bnlearn* [1], *shiny* [2], *pscl* [3], *dagitty* [4], *Future* [5], *visNetwork* [6], *bnviewer* [7], *igraph* [8], *DT* [9], *htmlwidgets* [10], *shinyscreenshot* [11], *shinythemes* [12], *shinyBS* (https://ebailey78.github.io/shinyBS), *tibble* (https://tibble.tidyverse.org/), *colourpicker* (https://github.com/daattali/colourpicker), *shinydashboard* (http://rstudio.github.io/shinydashboard/), *shinydashboardPlus* (https://github.com/RinteRface/shinydashboardPlus), *shinyjs* [13], *stringr* (https://stringr.tidyverse.org) and *purrr* (https://purrr.tidyverse.org/). For more details about the role of each R package in SAMBA, please refer to the Section “Material and Methods” of the main paper.

To start using SAMBA, BN models must be created and trained using the “*Build”* module that, via dropdown menu, splits in three Guide User Interfaces (GUI) named “*Learning and training*”, “*Load Network”*, “*Evidence/Control*”. As shown in Supplementary Figure S1, The BN model is created in the “*Learning and training”* GUI using two CSV or plain text files as input, one containing experimental variables data, and another including taxa raw counts (this dataset is available at the section “Data Availability Statement” of the main paper). The “*Learning and training”* interface presents two tab options called “*Experimental Variables Review”* and *“Count Data Review”* that permit the user to review the experimental variables and the abundance counts, respectively.

The tab “*Experimental Variables Review*” permits to visualize a summary with the experimental variable data. The tool works with discrete and continuous variables. Continuous experimental variables can be discretized in this section by selecting one or more variables via a dropdown menu containing variable’s names, and using one of the following methods: quantile, interval or Hartemink [14]. The last method (Hartemink) attempts to maximize mutual information between variables. If continuous variables are not discretized, a Shapiro test [15] is performed to know if these variables follow a normal distribution and if it is not the case, a log transformation is automatically performed for these variables. The raw counts for a given taxon in a sample are normalized by applying the following sum scale factor formula:

$$NC_{ij}=\frac{X_{ij}*\sum_{j=1}^{n} X_{ij}}{\left( \sum_{i=1}^{t} X_{ij} \right)}$$

where ${NC}_{ij}$ is the normalized count for a taxon (i) in a concrete sample (j), Xij is the raw counts of a concrete taxon (i) in a specific sample (j), n is the number of samples in the dataset and t is the number of taxa in the dataset. Note that the discretization of experimental variables will affect relationships that will be established during learning and training steps of the BN model building.

The tab “*Count Data Review*” allows the user to visualize raw and normalized abundance counts and edit or change taxa names to shorter ones. The user, in order to correct the inter-sample technical and/or biological variability among microbiomes of distinct specimens, can also apply here the zero-sum filter, to remove taxa with total zero normalized counts before building the BN model, and the prevalence filter, to remove taxa by their abundances or by their percentage in samples. This filter has two options: a) Total, where all samples are considered to remove taxa; b) Group, where samples are grouped by different states of an experimental variable indicated by the user. For each state, a percentage should be introduced and then, a taxon is removed if it does not fulfil the defined threshold. This filter can be applied after or before building the BN model.

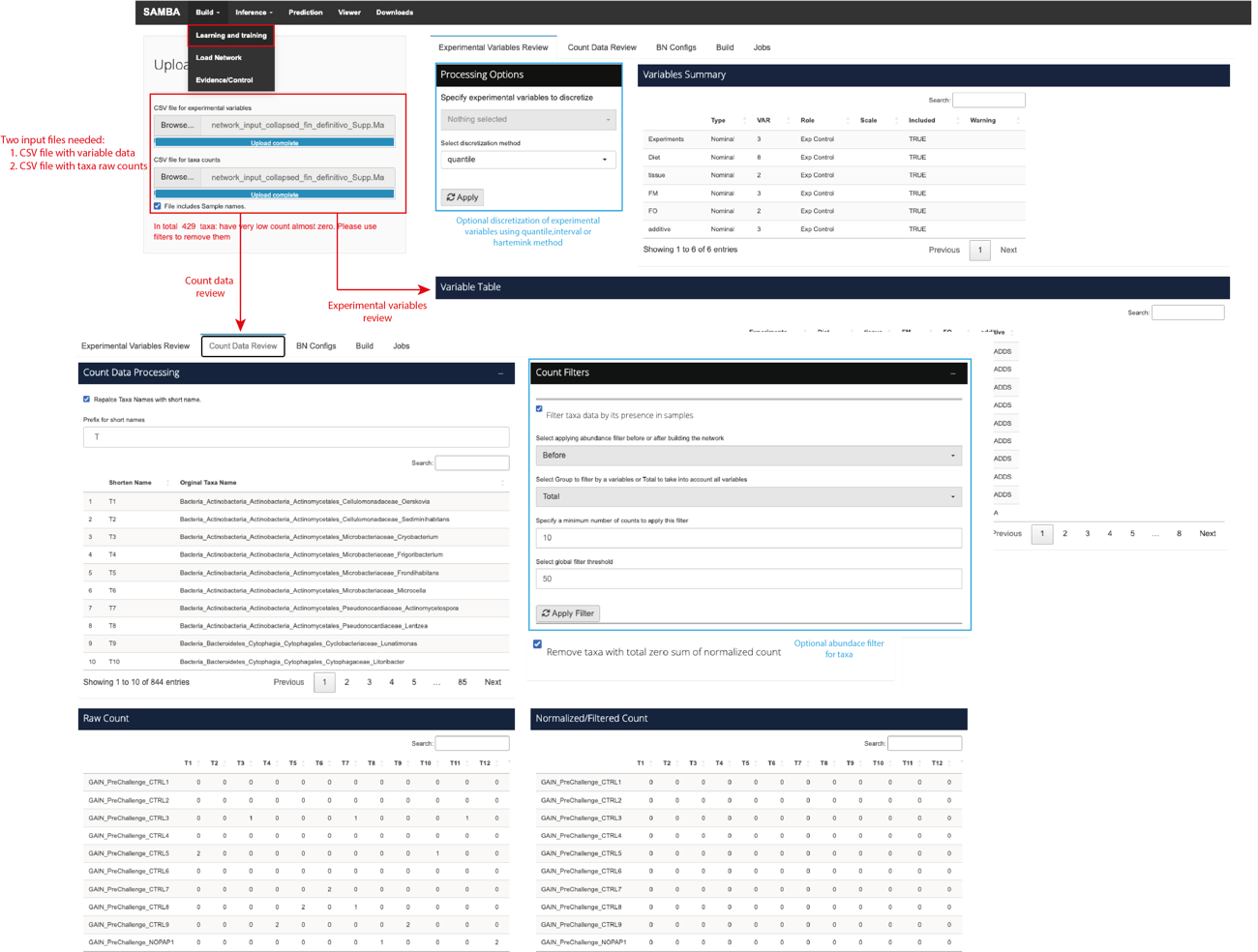

**Supplementary Figure S1**. Learning and training GUI. At the Top, interface for “Experimental Variables Review” tab, including a variable’s summary and a variable table, and including the option for discretizing one or more of them if needed. Below, the interface for “Count Data Review” tab, which includes an option to change taxa long names to shorter ones, an option to filter taxa by their abundances in samples, and two tables that show raw and normalize/filtered counts by taxon and sample.

After reviewing the experimental variables and the abundance counts, the user can modify or adapt the configuration options for building the BN model in the tab “*BN Configs*”. These options include: *“Network score”* (AIC, BIC or Loglik), *“Taxa distribution”* (Log-Normal or ZINB), *“Link strength: Mutual Information threshold”, “Link strength: Bayesian Information Criterion (BIC) threshold”*, *“Remove Experimental Variables interactions/relations from the network model structure”, “Blacklist”* and *“Whitelist”* (Supplementary Figure S2). The option “*Remove Experimental Variables interactions/relations from the network model structure”* allows the user to predict taxa values in a set of conditions not studied before in an experiment. Applying blacklists and/or whitelists, included in the hc() function of *bnlearn*, allow the user to consider or dismiss specific relationships within the input dataset. Regarding taxa distribution, if the Log-Normal Distribution is selected, the normalized count (NC) is log-transformed using the following formula: $(ln(NC+1) )$. Then the log-transform normalized data are merged to experimental variables data in one R object. For the ZINB distribution, normalized counts are combined with the experimental variables data without log-transformation.

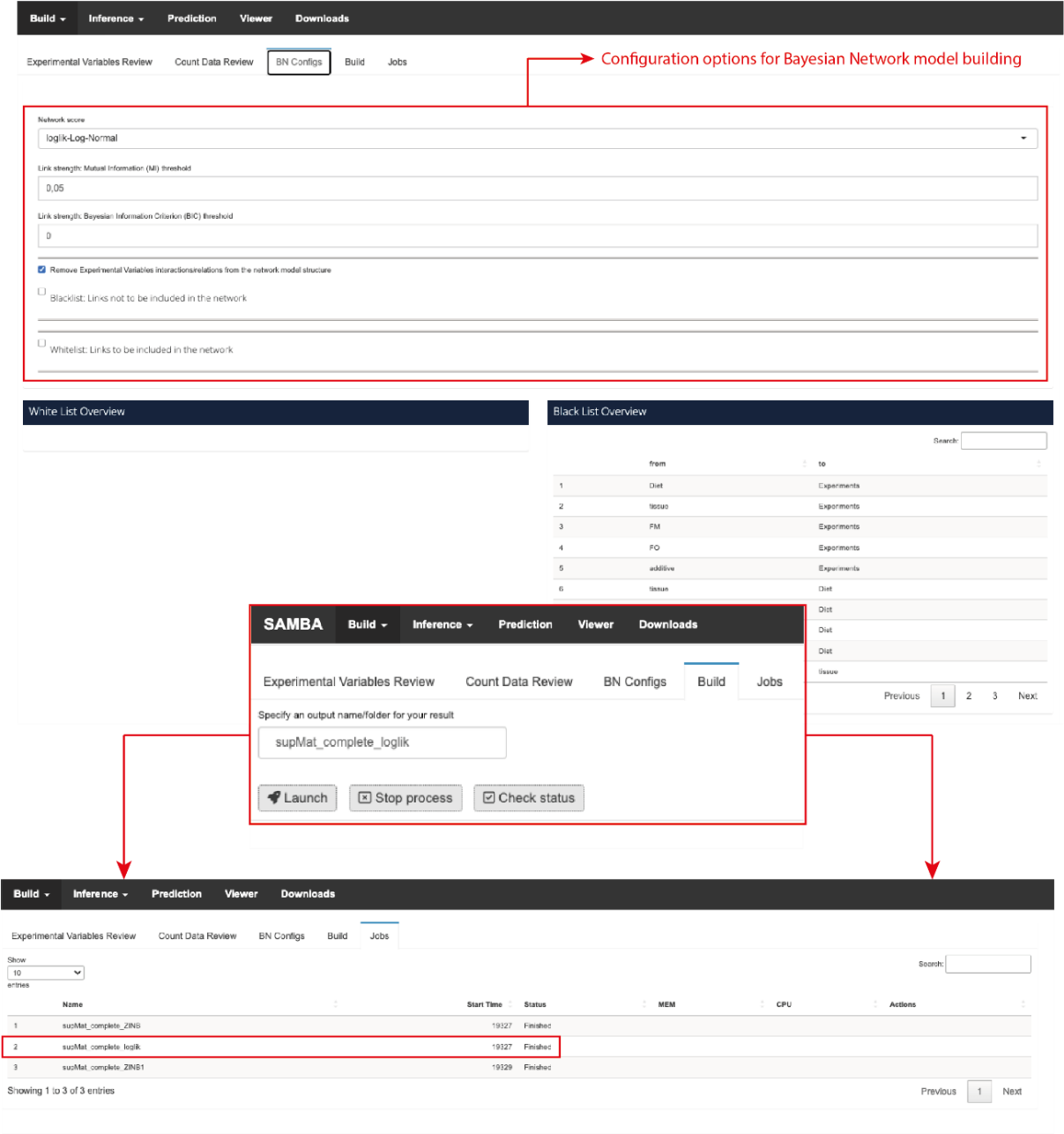

**Supplementary Figure S2**. Top, interface to configure Bayesian network model building, including network score, taxa distribution, link strength thresholds (Mutual Information and BIC), blacklist, whitelist and *“Remove Experimental Variables interactions/relations from the network model structure”* option. Also, a table for the whitelist overview and a table for the blacklist overview are shown. Middle, interface for building the BN model. Below, interface for tracking current or past launched jobs.

When the BN model is created, the user can download a zip file with the results (mainly the RData file) via the *“Download”* module (Supplementary Figure S3). Outputs of this BN model include an *RData* file containing the BN relationships modelled among variables and taxa; two text files containing the strength values of each link using BIC and MI criterions; two CSV file containing normalized and log normalized taxa counts; a log file; two CSV files including log normalized counts of taxa that passed the introduced conditions, and taxa that have been removed from the network.

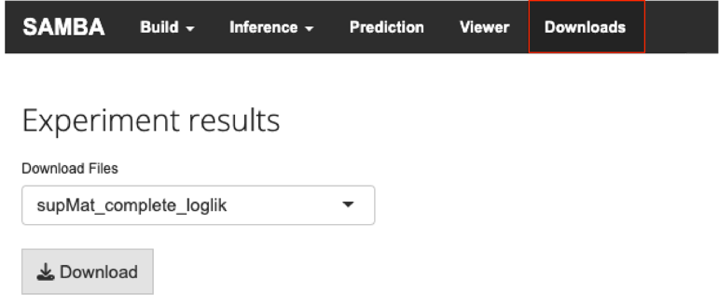

**Supplementary Figure S3**. Interface to download the BN model from the *“Download”* module.

Then, for further analysis in SAMBA, the user must upload the *RData* file including the BN model in the *“Load Network”* GUI of the “*Build*” module as shown in Supplementary Figure S4 (an example of .RData file is available at the link provided in the section “Data Availability Statement” of the main paper).

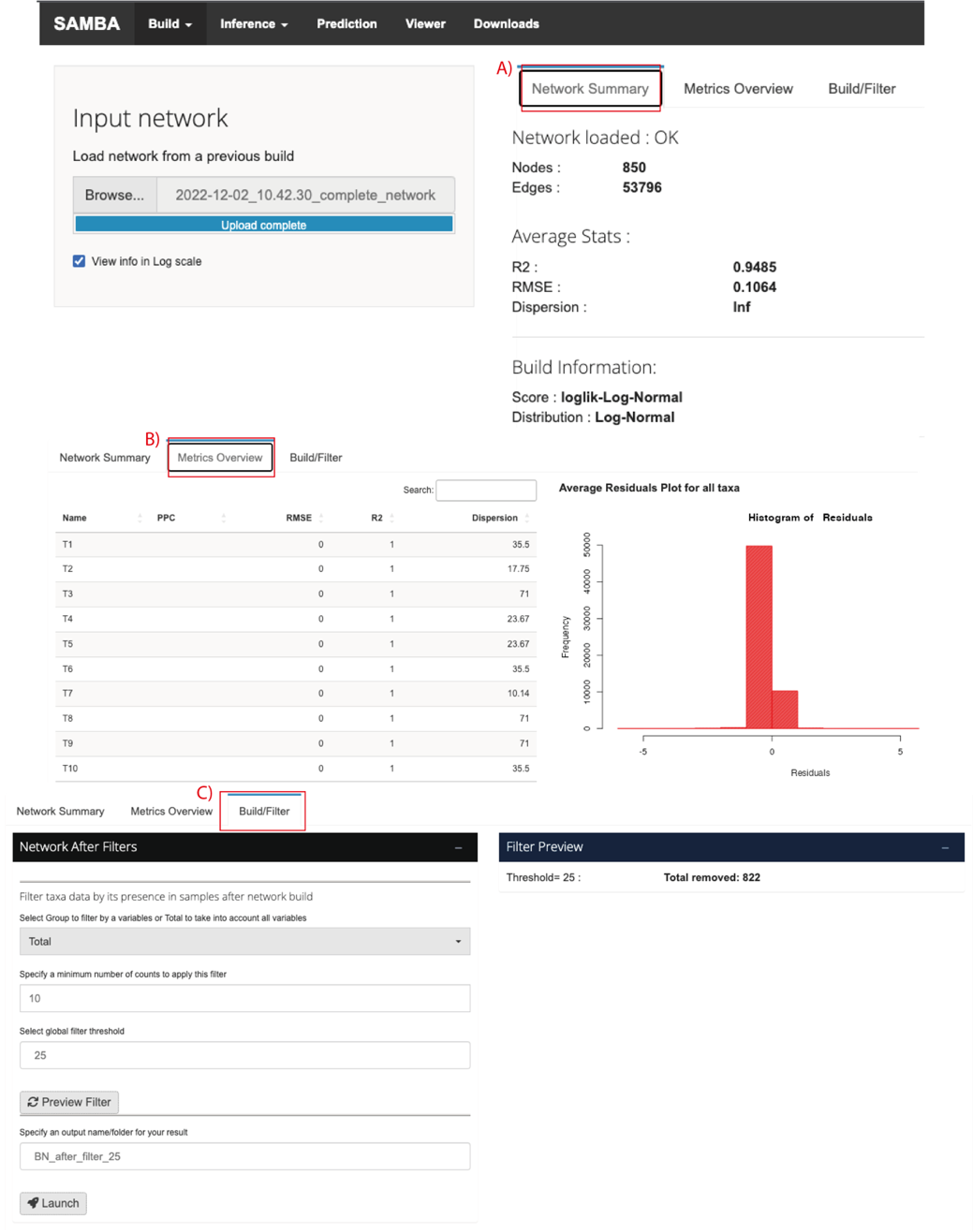

**Supplementary Figure S4**. Interface to upload the generated BN model to SAMBA. A) “*Network Summary”* option including basic information about the network. B) *“Metrics Overview”* option including statistics about taxa and a histogram for the average residuals for all taxa. C) “*Build/Filter*” option including the prevalence filter to be applied after building a BN model.

The “*Load Network*” GUI also gives basic information about the created BN model in the *“Network Summary”* tab option, including the number of nodes and edges of the model, average statistics and building information. In addition, the *“Metrics Overview”* tab displays a table containing information about Posterior Predictive Check (PPC), Root Mean Square Error (RMSE), R^2^ and dispersion of taxa; and a histogram showing the average residuals for all taxa. Finally, the “*Build/Filter*” tab allows the user to use the prevalence filter after the BN model has been built, removing those taxa whose raw counts are under a specific value in a percentage of samples.

Once the *RData* file is uploaded, the user can display and compare how data is distributed in different conditions or evidences by using the *“Evidence/Control”* GUI of the “*Build*” tab, as shown in Supplementary Figure S5. Here, by clicking on a taxon name, the following information is displayed:

1) a table for a combination including the number of samples that match the evidence, the mean value, the standard deviation and quartiles for each taxon (Supplementary Figure S5C).
2) a histogram for reference and target combination showing the frequencies of the normalized counts (Supplementary Figure S5D).
3) a density plot showing kernel density estimate, a smoothed version of the histogram (Supplementary Figure S5B).

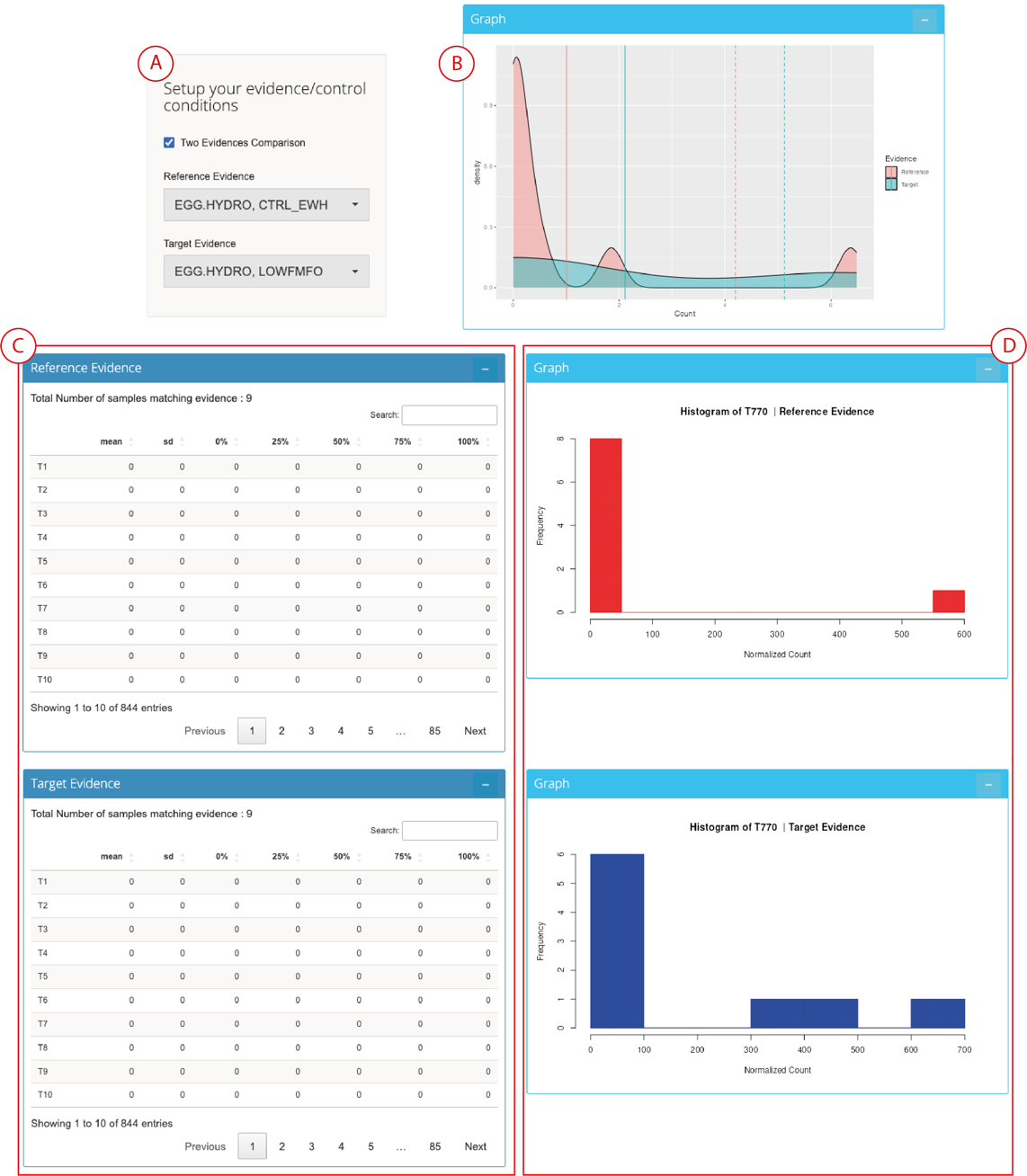

**Supplementary Figure S5**. *“Evidence/Control”* GUI. A) Global settings of the comparison. In this example, conditions EGG.HYDRO and CTRL_EWH (reference) are compared against EGG.HYDRO and LOWFMFO (target). B) Density plot of the comparison. The reference is colored in red and the target in blue. C) Tables with information about taxa in reference and target conditions. D) Histogram plotting frequencies of taxon T770 normalized counts in reference (red) and target (blue) conditions.

Next, SAMBA can infer or predict information from the aquaculture system interrogating the BN model using the *“Inference”* and *“Prediction”* modules. In this regard, the *“Inference”* module permits the user to infer information about which and how specific farming factors influence and shape the diversity of a given aquaculture pan-microbiome. To this end, the user only needs to upload the *RData* in the *“Load Network”* GUI of the *“Build”* module and, then, go to the “*Inference”* module. Within this module the user can get reports under two different options (“CPTs” or “DAG”) for both continuous and discrete variables. Under the “CPTs” option and for continuous variables (either taxa or farming factors), SAMBA displays CPTs and information about their relationships (negative or positive) with other taxa or farming factors in different discrete parent configurations plus a histogram of the residuals and a normal Q-Q plot (Supplementary Figure S6).

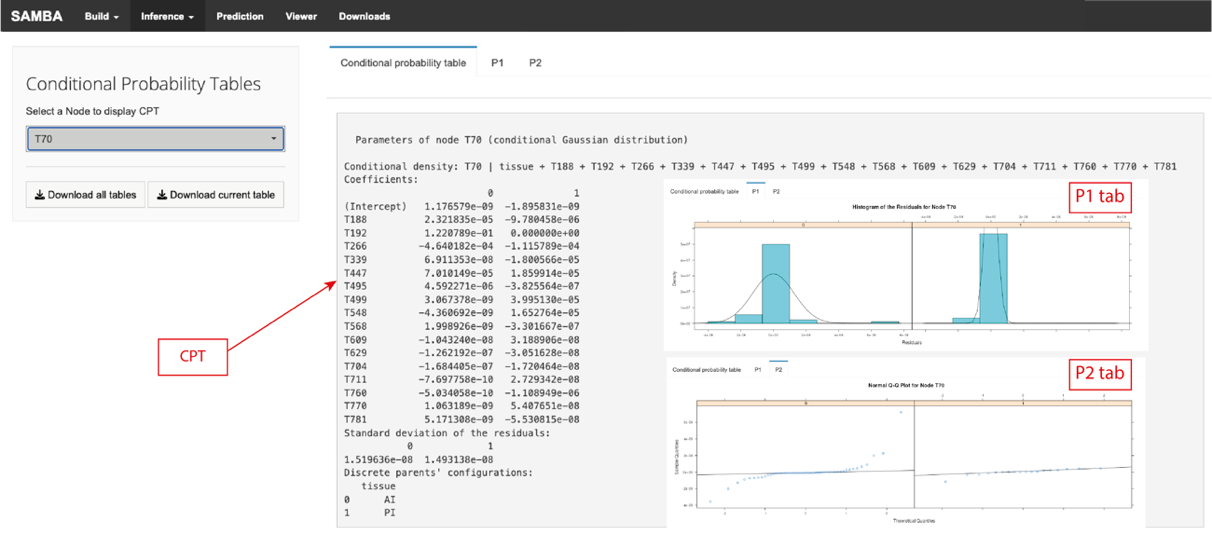

**Supplementary Figure S6**. CPT, histogram of residuals and normal Q-Q plot for a given taxon.

For discrete variables, the “CPTs” option also displays the CPT information including the probability of each state given other discrete variables and a plot of conditional probabilities for the selected node (Supplementary Figure S7).

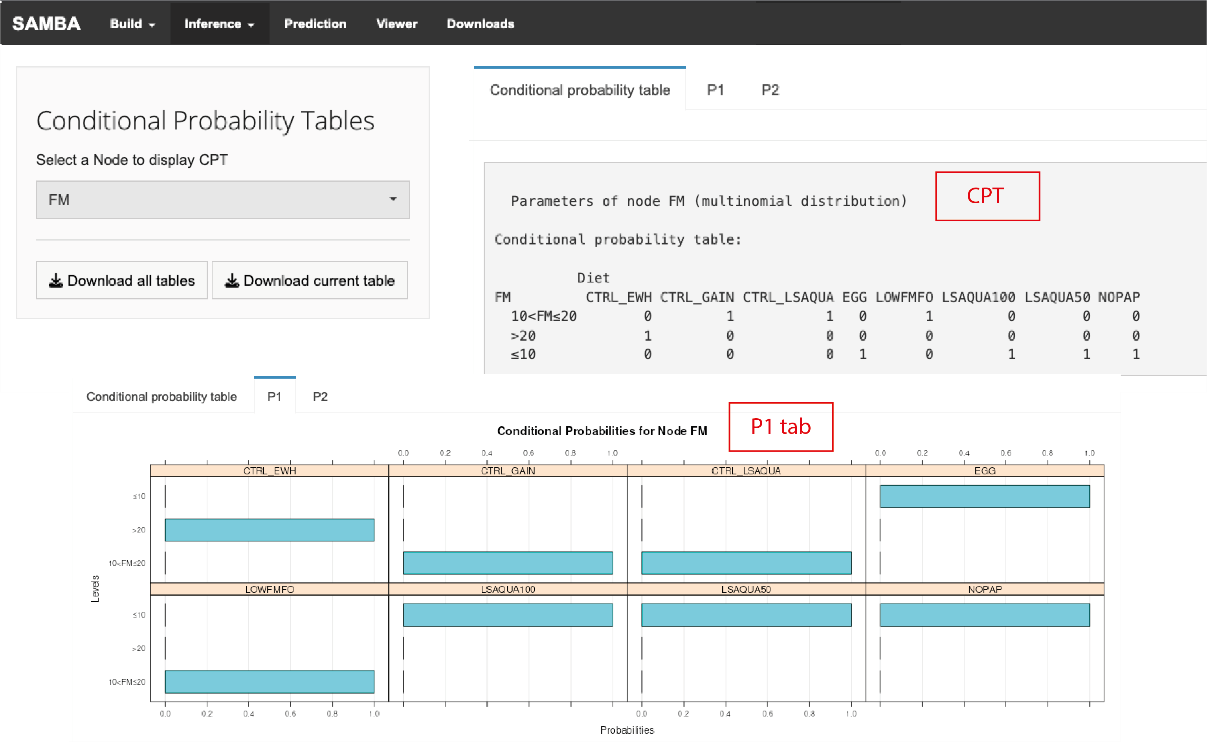

**Supplementary Figure S7**. CPT and conditional probabilities plot for the FM node.

In the “DAG” option, direct and/or indirect relationships of a given taxon node (Markov blanket) are displayed (Supplementary Figure S8). Conditional dependency with experimental variables and testable implications about conditional independence are also displayed in this option.

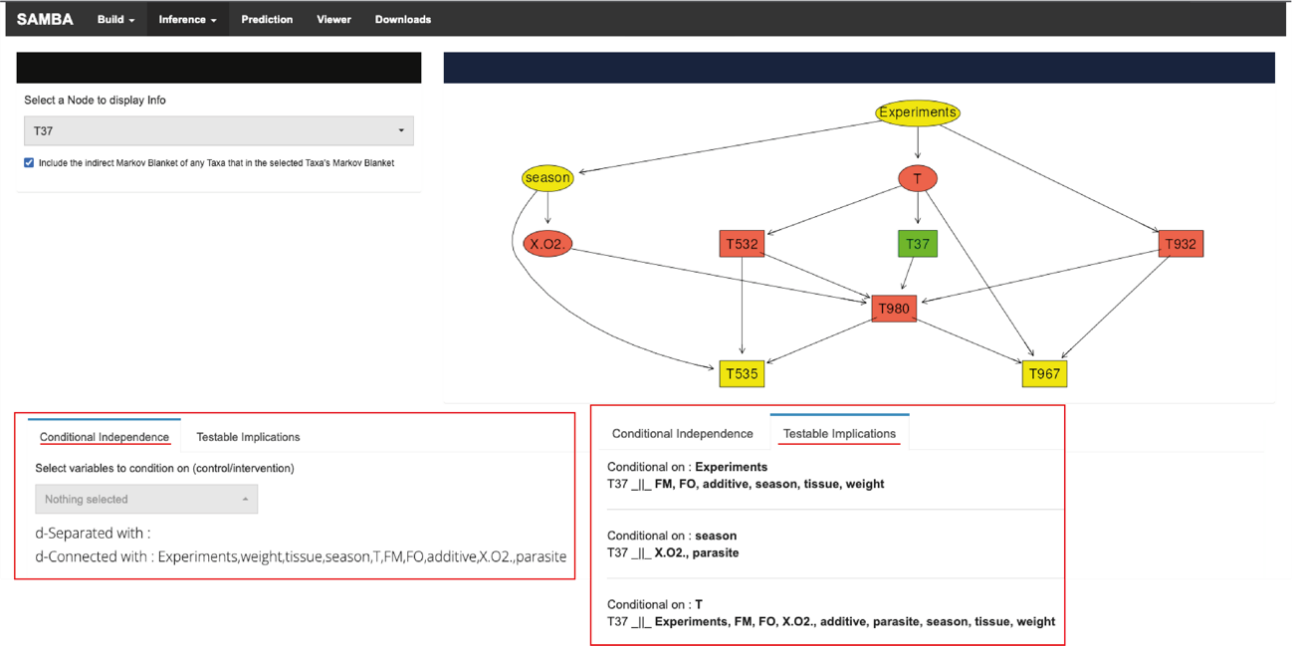

**Supplementary Figure S8**. DAG option interface for the node T37, highlighted green. Nodes with an indirect relationship are colored in yellow and nodes with a direct relationship are colored in red. Nodes corresponding to taxa are represented by a square shape while nodes corresponding to experimental variables are represented by an oval shape. Conditional independence and testable implications for node T37 can also be displayed.

The “*Prediction”* module provides two pipeline options: “*Predict abundances*” and “*Predict Metagenomes”*. The first option allows the user to get a predicted range of abundances for one or more taxa given specific experimental conditions, while the second option can be used to infer the metagenomic profile from a specific microbiome profile. As shown in Supplementary Figure S9A, to perform the prediction of abundances, users must upload the *RData file* with the BN model in the “*Load Network*” interface. Once the model has been introduced in SAMBA, users must acess the tab *“Predict Abundances”* to select the taxa names, of which the user wants to predict the abundance counts, and to select the states or conditions of the variables as conditional evidence. In the Log-Normal distribution mode, normalized frequencies in log scale are obtained via the cpdist() function of *bnlearn*. This function returns a data frame containing the samples generated from the conditional distribution of the nodes according to the states selected for each experimental discrete variable in a combination and weight of each value. In ZINB distribution mode, custom sampling method was implemented to sample from the fitted ZINB models of each taxon in the BN. First, a sampling path is extracted from the network based on node hierarchy. Then, the sampling path is used to sample each node of the path in order by random sampling from a ZINB distribution using fitted model coefficients. To account for uncertainty propagation through the sampling path, the expected mean of the model is reported beside the sample's means.

Predictions are then shown as an exportable table in the GUI (Supplementary Figure S9B). The module infers the following metrics for predicted and observed abundance counts: a) Pred. Average: Expected/average of the normalized count; b) Pred. SD: The standard deviation of the predicted count in all samples; c) Pred. Q-0%; Pred. Q-25%; Pred. Q-50%; Pred. Q-75%; Pred. Q-100% columns: Quartile ranges; d) Pred. M+SD: Deviation of the predictions around the mean range [mean ± standard deviation]; e) Pred. P(M+SD) column: Probability density value of the range, which is inferred from the probability density of the generated samples, range from 0 to 1; f) Pred. HDR_50%: Range of the Highest Density Region, which is the region having the smallest possible range that covers the sample space for a 50% of probability; g) Obs. Data columns: Rest of columns in the output providing Inferences obtained from the original input data using the same evidence, including average normalized counts for each taxon based on the same evidence using mean() function and the sd() function implemented in R. A correction step is implemented in this pipeline for applying prevalence filters or for adjusting the prediction (using the input data as a prior) to re-infer corrected posterior samples until the prediction converges with real-world observation. In particular, using the Bayes rule, the probability density of the generated samples is multiplied by the probability density of the input data given the input evidence to get new posterior samples. Then, a weighted average of the normalized predicted value (NPV) is calculated for the generated samples using the following formula:

$$NPV=e^{\left( \frac{\sum_{i=1}^{n} P_{i}*W_{i}}{\sum_{i=1}^{n} W_{i}} \right)}-1$$

where “n” is the number of samples generated from the network, “i” is a specific generated sample, “P” is the value of the generated sample and W is the weights of a generated sample.

In the “Results and discussion” of the main paper, we present and discuss the results obtained testing the “*Predict abundances*” function of SAMBA with two empirical datasets (available at the Section “Data Availability Statement” of the main paper).

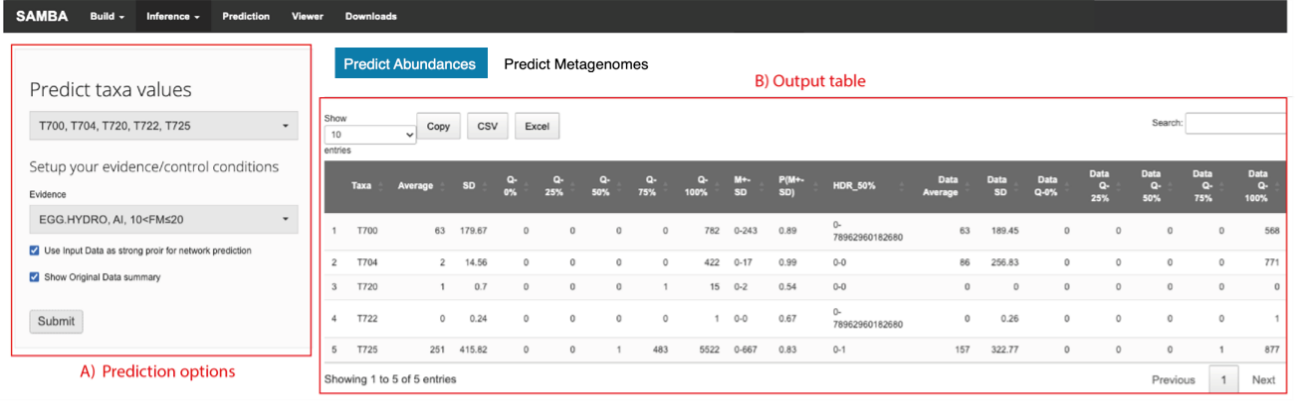

**Supplementary Figure S9**. GUI for predicting abundance values of some or all taxa of the microbiome depending on the state or conditions of other variables. A) Prediction options to select taxa and experimental conditions (Evidence). B) Output table containing prediction data and original data summary.

As mentioned above, the *“Prediction”* module also permits the users to predict the metagenomic profile of a given microbiome using the pipeline available via the *“Predict Metagenome”* tab provided by PICRUSt2 [16]*.* As input, the users needs to upload a text file containing the taxa abundances in samples, and a FASTA file including taxa reference sequences. PICRUSt2 protocols include a KEGG and a MetaCyc option to get information about KEGG [17] or MetaCyc [18] pathways of the predicted metagenomic profile, respectively. Independently of the protocol selected, PICRUSt2 performs four steps:

i. Place sequences into a reference tree based on 20,000 16S rRNA sequences from genomes in the Integrated Microbial Genomes database [19]. This step consists of aligning taxa sequences with a multiple-sequence alignment of reference 16S rRNA sequences with HMMER [20]; Finding the most likely placements of the taxa in the reference tree with EPA-NG [21] or SEPP [22]; Outputs of a treefile with the most likely placement for each taxon with GAPPA [23]. The output file is called out.tree.
ii. Hidden-state prediction of gene families, that predicts the copy number and enzymes of gene families for each taxon. Two gz files are generated as output: 16S_predicted_and_nsti.tsv.gz, containing predicted 16S copy numbers for all study sequences, and EC_predicted.tsv.gz if MetaCyc mode is selected, containing predicted Enzyme Commission (EC) number abundances for all sequences, or KO_predicted.tsv.gz if KEGG option is selected, containing predicted KEGG Orthology (KO) abundances for all sequences.
iii. Generate metagenome predictions. Per-sample metagenome functional profiles are generated based on the predicted functions for each study sequence. The specified sequence abundance table will be normalized by the predicted number of marker gene copies. The prediction generates five output files: a) pred_metagenome_unstrat.tsv.gz: overall EC number or KO abundances per sample and taxon; b) pred_metagenome_unstrat_descrip.tsv.gz: functional descriptions added to pred_metagenome_unstrat.tsv.gz file; c) pred_metagenome_contrib.tsv.gz: a stratified table in “contribution” format breaking down how the taxa contribute to gene family abundances in each sample; d) seqtab_norm.tsv.gz: the taxa abundance table normalized by predicted 16S rRNA copy number; e) weighted_nsti.tsv.gz: the mean Nearest Sequenced Taxon Index (NSTI) value per sample considering the relative abundance of the taxa. This file can be useful for identifying outlier samples in the dataset.
iv. Pathway-level inference, which infers the presence and abundance of MetaCyc or KEGG pathways based on gene family abundances in a sample. There are several output files created within the pathways out directory: a) path_abun_contrib.tsv.gz: a stratified table in “contributional” format breaking down how the taxa contribute to pathway abundances in each sample; b) path_abun_unstrat.tsv.gz: overall pathway abundances per sample; c) path_abun_unstrat_descrip.tsv.gz: functional descriptions added to path_abun_unstrat.tsv.gz file.

Input file examples can be downloaded through the link provided in the Section “Data Availability Statement” of the main paper.

Finally, to view, custom, edit for publication or just investigate the BN model, the user must go to the “*Viewer*” module and click on the “Refresh” button to visualize the network structure. To use this module, users must load the RData file of the BN model into the “*Load Network*” interface of the “*Build*” module. As shown in Supplementary Figure S10, the “Viewer” module is composed by the following sections: A) global settings; B) a graph panel; C) “Edit”, “Select” and “Filter” menu; D) a data manager for nodes and edges, and E) node and edge information table.

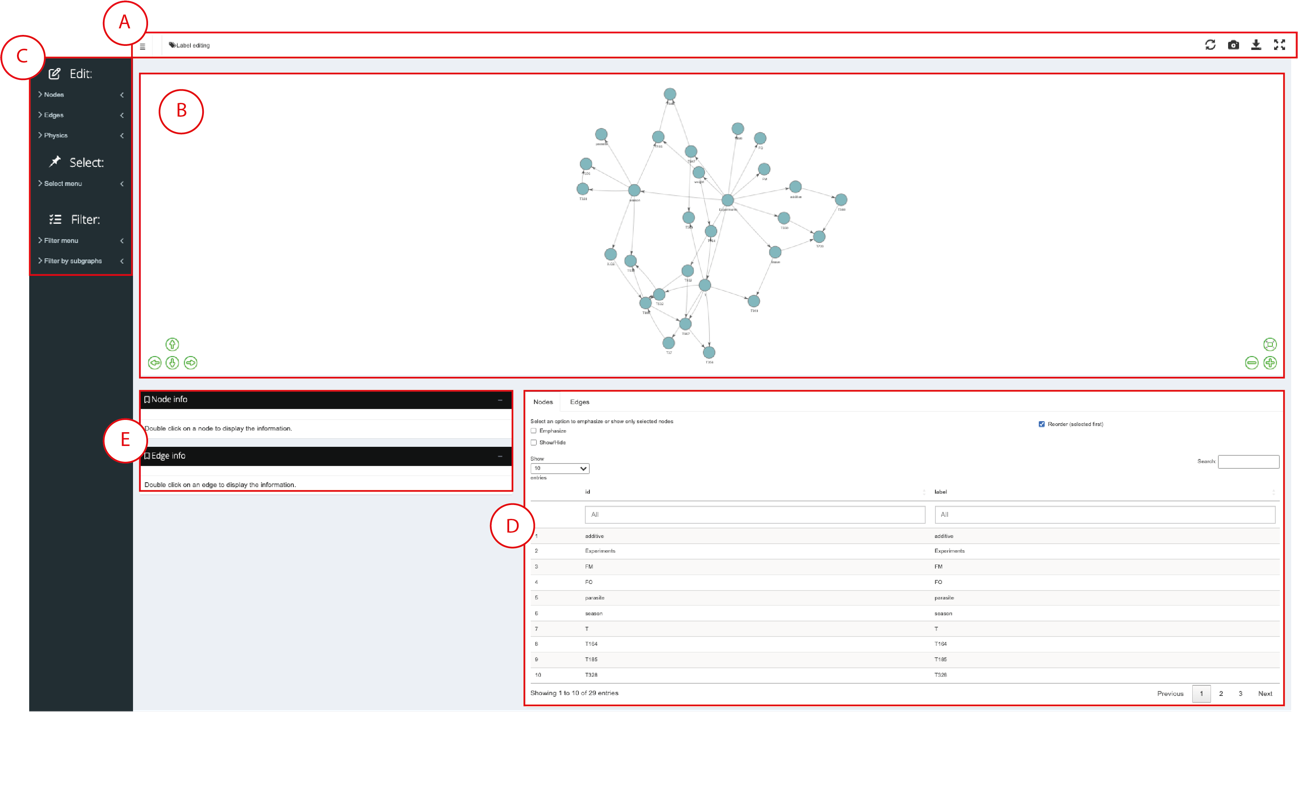

**Supplementary Figure S10**. Network Viewer interface. A) Global settings, including “Display the edition sidebar”, “Label editing”, “Refresh Network”, “Take a screenshot”, “Save network” and “Full-screen mode”. B) Graph panel including options to change the network’s position, readjust its size and position, and enlarge or reduce the size. C) Menu that contains options to modify nodes’ and edges’ shape, size, color, shadow and label (Edit menu); to select a subset of nodes related to the node that the user selected by their link’s directions and their grade (Select menu); and to create node groups by selecting them in the graph or by entering node names, and to emphasize or just show a particular subgraph (Filter menu). D) Data manager including information about nodes and edges in a table format. E) Node and Edge info showing node’s CPT and edge’s strength, respectively.

The graph panel allows the user to visualize the network and to change its position, enlarge or reduce its size and return to its original position. In addition, graph nodes can be moved at the user’s convenience by clicking on them and dragging them. The user can modify nodes’ and edges’ color, size, label characteristics, shape and shadow by clicking on nodes or edges of interest and changing these features in the “Edit menu”. This menu has also an option named “Physics”, which includes functions to fix X and/or Y axis, so the graph panel will be fixed, or to give a bounce effect to the network. Nodes can not only be selected manually, but can also be selected using the “Select” menu. If a node is chosen, the user can select, through this menu, a group of related nodes that are parents and/or children (“Direction” option) of this node given a certain grade of distance (in edges) among them. An additional option to create a set of nodes is to pick them in the “Nodes table”. When a set of nodes is selected, the users can create a group using the “Filter” menu to emphasize or to hide nodes that are not present in this set, so the users can work with the nodes they are interested in. An alternative to create a group in this menu is to write node names in the “Input a list of nodes” option or to select a specific subgraph of the network using the “Filter by subgraphs” option. When a set or a group of nodes is selected, nodes’ and edges’ feature modifications can be applied in all of them at the same time using the “Edit” menu. Once the network has been edited and/or filtered, the user can save it in several formats such as HTML, PDF, PNG or JPEG.

Furthermore, in this module, the user has information about the CPT of a selected node (“Node info” section) , the strength of a selected edge (“Edge info” section), a table containing the label and the identifier of nodes (“Nodes” tab), and a table including established edges between nodes, their identifiers and their strengths (“Edges” tab).

A full user guide for managing the *“Viewer”* module of SAMBA is available at the link provided in the Section “Data Availability Statement” of the main paper.

**2.- References**

1.	Scutari, M. Learning Bayesian Networks with the bnlearn R Package. *Journal of Statistical Software* **2010**, *35*, 1 - 22, doi:10.18637/jss.v035.i03.

2.	Chang, W.; Cheng, J.; Allaire, J.; Stievert, C.; Schloerke, B.; Xie, Y.; Allen, J.; McPherson, J.; Dipert, A.; Borges, B. shiny: web application framework for r. R package version 1.7.4. Available online: (accessed on 23 June 2023).

3.	Zeileis, A.; Kleiber, C.; Jackman, S. Regression Models for Count Data in R. *Journal of Statistical Software* **2008**, *27*, 1 - 25, doi:10.18637/jss.v027.i08.

4.	Textor, J.; van der Zander, B.; Gilthorpe, M.S.; Liskiewicz, M.; Ellison, G.T. Robust causal inference using directed acyclic graphs: the R package 'dagitty'. *Int J Epidemiol* **2016**, *45*, 1887-1894, doi:10.1093/ije/dyw341.

5.	Bengtsson, H. A Unifying Framework for Parallel and Distributes Processing in R using Futures. *The R Journal* **2021**, *13*, 273-291.

6.	Almende, B.; Thieurmel, B.; Robert, T. visNetwork: Network Visualization using’vis. js’ Library. R package version 2.0.9. Available online: (accessed on 23 June 2023).

7.	Fernandes, R. bnviewer: Bayesian networks interactive visualization and explainable artificial intelligence. R package version 0.1.6. Available online: (accessed on 23 June 2023).

8.	Csardi, G.; Nepusz, T. The igraph software package for complex network research. *InterJournal, complex systems* **2006**, *1695*, 1-9.

9.	Xie, Y.; Cheng, J.; Tan, X. DT: A Wrapper of the JavaScript Library 'DataTables'. R package version 0.26. Available online: (accessed on 23 June 2023).

10.	Vaidyanathan, R.; Xie, Y.; Allaire, J.J.; Cheng, J.; Sievert, C.; Russell, K. htmlwidgets: HTML Widgets for R. R package version 1.6.0. Available online: (accessed on 23 June 2023).

11.	Attali, D.; von Hertzen, N.; Grey, E. shinyscreenshot: Capture Screenshots of Entire Pages or Parts of Pages in 'Shiny'. R package version 0.2.0. Available online: (accessed on 23 June 2023).

12.	Chang, W. shinythemes: shinythemes: Themes for Shiny. R package version 1.2.0. Available online: (accessed on 23 June 2023).

13.	Attali, D. shinyjs: Easily Improve the User Experience of Your Shiny Apps in Seconds. R package version 2.1.0. Available online: (accessed on 23 June 2023).

14.	Hartemink, A.J. Principled computational methods for the validation discovery of genetic regulatory networks. Massachusetts Institute of Technology, 2001.
